# PharaCon: A new framework for identifying bacteriophages via conditional representation learning

**DOI:** 10.1101/2024.06.16.599237

**Authors:** Zeheng Bai, Yao-zhong Zhang, Yuxuan Pang, Seiya Imoto

**Affiliations:** Division of Health Medical Intelligence, Human Genome Center, The Institute of Medical Science, The University of Tokyo, Minato-ku, Tokyo, Japan; Collaborative Research Institute for Innovative Microbiology, The University of Tokyo, Bunkyo-ku, Tokyo, Japan

## Abstract

**Motivation:** Identifying bacteriophages (phages) within metagenomic sequences is essential for understanding microbial community dynamics. Transformer-based foundation models have been successfully employed to address various biological challenges. However, these models are typically pre-trained with self-supervised tasks that do not consider label variance in the pre-training data. This presents a challenge for phage identification as pre-training on mixed bacterial and phage data may lead to information bias due to the imbalance between bacterial and phage samples.

**Results:** To overcome this limitation, this study proposed a novel conditional BERT framework that incorporates labels during pre-training. We developed an approach using a conditional BERT model for pre-training labeled data, incorporating label constraints with modified language modeling tasks. This approach allows the model to acquire label-conditional sequence representations. Additionally, we proposed a solution that utilizes conditional BERT in the fine-tuning phase as a classifier. We applied this conditional BERT framework to identify phages using a novel fine-tuning strategy, introducing PharaCon. We evaluated PharaCon against several existing methods on both simulated sequence datasets and real metagenomic contig datasets. The results demonstrate PharaCon's potential as an effective and efficient method for phage identification, highlighting the effectiveness of conditional B ERT as a solution for learning label-specific representations during pre-training on mixed sequence data.

**Availability:** The codes of PharaCon are now available in: https://github.com/Celestial-Bai/PharaCon.

## 1 Introduction

Bacteriophages (short as phages), known as bacterial viruses, play a crucial role in microbial communities. Identifying phages from mixed metagenomic sequences is often the first step in phage analysis. Traditional sequence alignment-based methods have been widely used in real-world applications, such as VIBRANT (Kieft *et al*., 2020) and VirSorter2 (Guo *et al*., 2021). These alignment-based methods are relatively computationally intensive, as they frequently align target sequences with existing profile hidden Markov models (Eddy, 1998).

With the advancement of deep learning technologies, learning-based methods have been proposed to address this limitation. State-of-the-art methods include the CNN-based DeepVirFinder (Ren *et al*., 2020) and the LSTM-based Seeker (Auslander *et al*., 2020). Recently, MetaPhaPred (Ma *et al*., 2023) utilized the attention mechanism and achieved top performance in their proposed experiments.

Leveraging the success of Transformers (Vaswani *et al*., 2017) in natural language processing (NLP) (Devlin *et al*., 2019) and computer vision (CV) (Dosovitskiy *et al*., 2020), Transformer-based foundation models using a pre-train-fine-tune paradigm have recently been applied to tackle biological challenges. Nucleotide sequence encoders like DNABERT (Ji *et al*., 2021), DNABERT-2 (Zhou *et al*., 2023), and Nucleotide Transformer (Dalla-Torre *et al*., 2023) have demonstrated their capabilities in addressing various genomic problems. These foundation models are used for learning generalized genomic features. During pre-training, these methods use self-supervised learning tasks without considering label variance. Even when the label information of pre-training data is known, it is not incorporated into the pre-training process, posing a challenge for pre-training a Transformer-based model for phage identification. Since the amount of known bacteria and phages is imbalanced, pre-training with mixed bacteria and phage data may lead to information bias in the model.

Previously, to address this potential risk, we proposed a method for phage identification that utilized label information in the pre-training data by pre-training bacterial and phage sequences in two separate DNABERT models and fine-tuning them simultaneously within a unified framework. This novel representation learning method, named INHERIT (Bai *et al*., 2022), achieved top performance according to our experimental results. INHERIT utilizes label information to some extent by splitting the imbalanced data into two groups to pre-train separately according to the labels. In this paper, we focus on a more efficient way to train a model that can learn label-specific features.

In this study, we proposed a novel architecture that can learn sequence representations with their labels. Previously, in natural language processing, Wu et al. (Wu *et al*., 2019) proposed a conditional BERT for data augmentation. The pre-trained conditional BERT model generates augmented data, which helps re-train traditional sequence classifiers for downstream tasks. The conditional BERT is not incorporated into the downstream tasks.

Different from their work, our proposed architecture shares the same name but introduces different and novel features:

1. We introduce the label classes as special tokens. Our conditional BERT model attaches the labels during tokenization. While in their work, label information is input as an embedding layer.
2. Instead of using conditional BERT for generating augmented data without utilizing the learned representations in the downstream tasks, we proposed a novel solution that applies conditional BERT to both pre-training and fine-tuning processes. We introduced a novel finetuning scheme that generates “paradox” samples during tokenization, making conditional BERT feasible for classification tasks. Not only do we pre-train the conditional BERT to learn label-specific contextual information, but we also directly fine-tune the conditional BERT as a classifier, predicting the same label classes used in pre-training.

We applied this novel architecture to phage identification task, and named this framework PharaCon. Based on our experimental results, PharaCon achieved top performance compared to selected existing methods in our simulated sequence dataset and was the best learning-based method in our real metagenomic contig dataset. It also proved to be more efficient than alignment-based methods. The source code for PharaCon can be found at https://github.com/Celestial-Bai/PharaCon.

## 2 Methods

### 2.1 Conditional BERT

#### 2.1.1 Label attachment and tokenization

To pre-train bacterial and phage samples with their own label constraints, we designed a BERT-style model called conditional BERT. We attached the corresponding label when tokenizing a metagenomic nucleotide sequence. Specifically, we adapted two special label class tokens: [BAC] and [PHA], which are abbreviations for bacteria and phages, respectively. If a tokenized sequence belongs to bacteria, we attach [BAC] at the beginning of the tokens; if a tokenized sequence belongs to phages, we attach [PHA].

To tokenize the nucleotide sequences, we utilized the overlapped k-mer tokenizer, following the settings of DNABERT (Ji *et al*., 2021). Although various tokenizers for nucleotide sequences with different strategies have been proposed recently, there is no definitive conclusion as to whether they are better suited to DNA sequences than the overlapped k-mer tokenizer (Marin *et al*., 2023). Moreover, to ensure a fair comparison with INHERIT in our experiments, we ultimately decided to use the overlapped k-mer tokenizer.

Figure 1 illustrates two examples of tokenization with DNABERT and our conditional BERT. For DNABERT, the class token [CLS] is attached regardless of whether the nucleotide segment belongs to a bacterium or a phage. Consequently, the model learns contextual information from a mixture of bacterial and phage samples without any constraints, which may cause information bias if bacterial and phage samples are imbalanced. However, conditional BERT attaches the label class tokens [BAC] and [PHA] during tokenization, instead of the traditional class token [CLS].

**Fig. 1:**
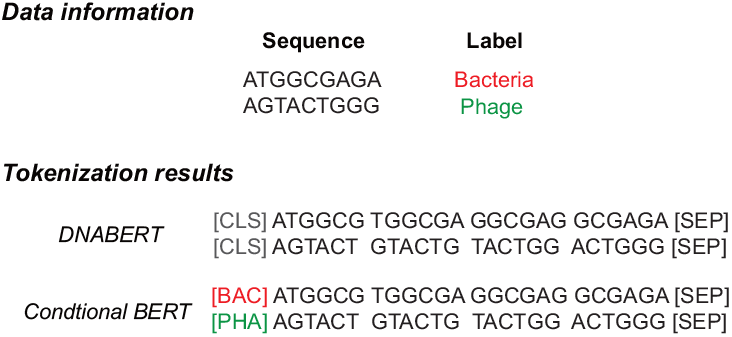
Examples of tokenization differences between DNABERT and our designed conditional BERT. DNABERT attaches label token [CLS] to every sample regradless of its label class. In contrast, conditional BERT attaches label tokens according to the label class to apply constraints to the pre-training process, allowing the model to learn the contextual information according to the label classes.

#### 2.1.2 Pre-training

Figure 2A illustrates the pre-training process of conditional BERT. Each 500 bp segment in our pre-training set is tokenized into overlapped k-mers (with *k* = 6 in our study). A separator token [SEP] is appended at the end, and a label token is attached at the beginning simultaneously. Following the settings of the traditional masked language modeling task (Devlin *et al*., 2019; Ji *et al*., 2021), during pre-training, we randomly masked 15% of the non-special tokens (label tokens and the separator token are considered special tokens). Among the masked tokens, 80% are replaced with the mask token [MASK], 10% are replaced with a token randomly selected from the vocabulary, and the remaining 10% are left unchanged. The model is trained to predict these masked tokens in each input sequence.

**Fig. 2:**
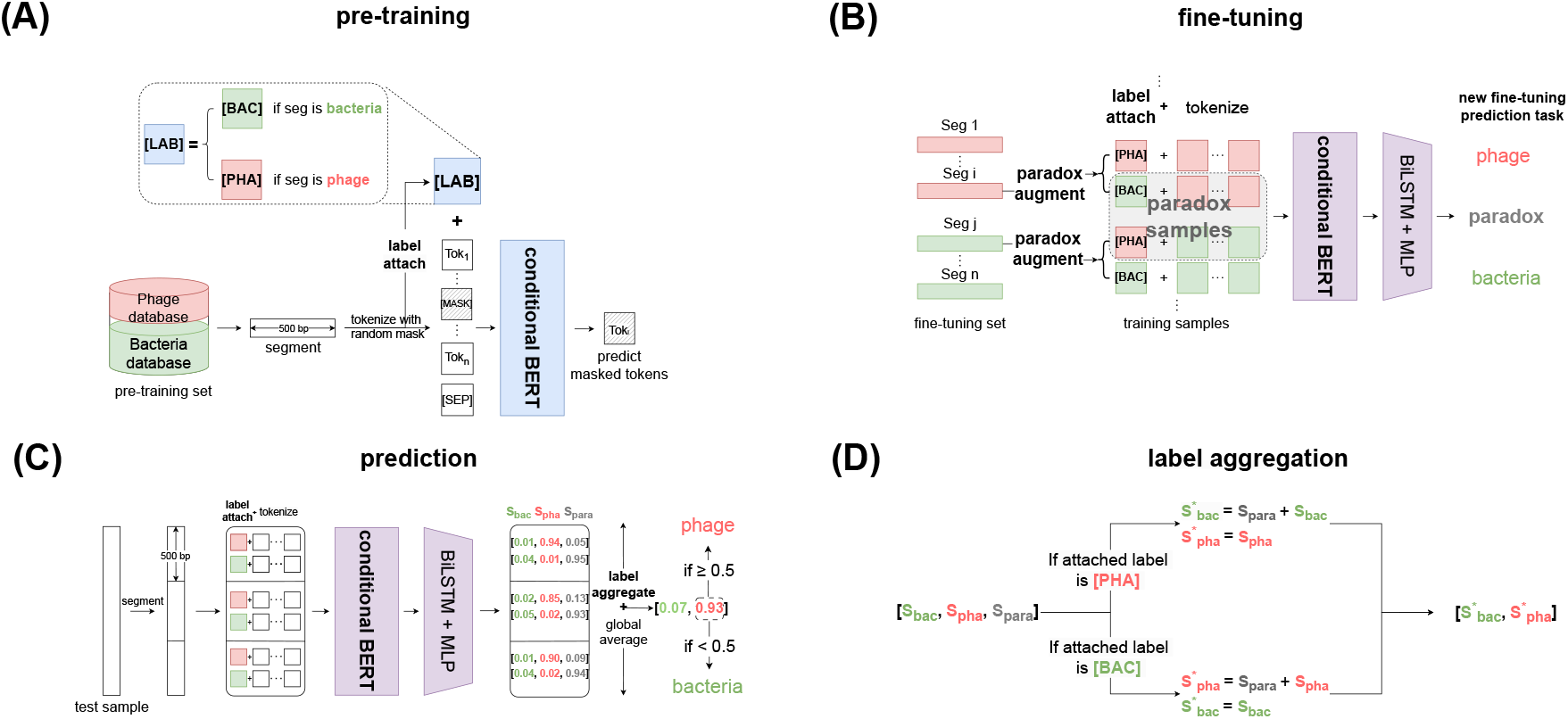
The pipeline of PharaCon. (A) Conditional BERT is pre-trained using labeled data with label tokens (e.g., [BAC], [PHA]) attached during tokenization. This allows learning label-specific contextual representations via masked language modeling. (B) During fine-tuning, sequences are tokenized twice with different label tokens, creating “paradox” samples with incorrect labels. The model is trained on these augmented samples to predict the correct label and identify paradoxes. (C) For prediction, metagenomic sequences are split into 500 bp segments, each tokenized with [BAC] and [PHA] labels. PharaCon generates label class scores, using a label aggregation strategy to convert three-label scores (bacteria, phage, paradox) into binary scores. These scores are averaged across segments for final classification. (D) The label aggregation strategy combines the scores. Scores for bacteria and paradox are combined when tokenized with the [PHA] label, and scores for phage and paradox are combined when tokenized with the [BAC] label.

To estimate a masked token in a sequence, the model predicts the probability distribution of tokens at the masked position, conditioned on the label constraint and the unmasked context of the sequence. The masked language modeling task can be summarized by the following formula:

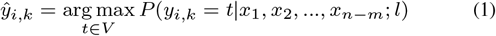

where 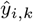 is the estimate of the *k*-th token at the masked position *i. V* is the set of unique tokens in the vocabulary and *t* is a token in *V* . *n* is the total number of tokens and *m* is the number of masked tokens. The model estimates the masked token, given the condition of (*n* − *m*) unmasked tokens {*x*_1_, *x*_2_, …, *x*_*n*−*m*_} and label constraint *l*.

#### 2.1.3 Label class representations are learned during pre-training

Pre-training with attached label class tokens enables the model to learn contextual information conditioned on specific labels. The label token representations differ after pre-training, demonstrating the model's ability to distinguish between different types of sequences. To validate this, we used the same pre-training bacterial and phage samples for pre-training a traditional DNABERT model. The detailed pre-training dataset information and settings will be described in Section 3.1.

We randomly sampled 10,000 bacterial segments and phage segments from the pre-training set and generated the class token representations (the last hidden state of the class tokens) using both conditional BERT and DNABERT. We then visualized these representations, combined with the word embeddings of the class tokens, using t-SNE plots (see Figure 3). The t-SNE plot for DNABERT shows that the class token representations are mixed together, and indicating that the contextual information learned by DNABERT does not differentiate well between bacterial and phage sequences. This is because DNABERT uses the same class token regardless of whether the sample is bacterial or phage.

**Fig. 3:**
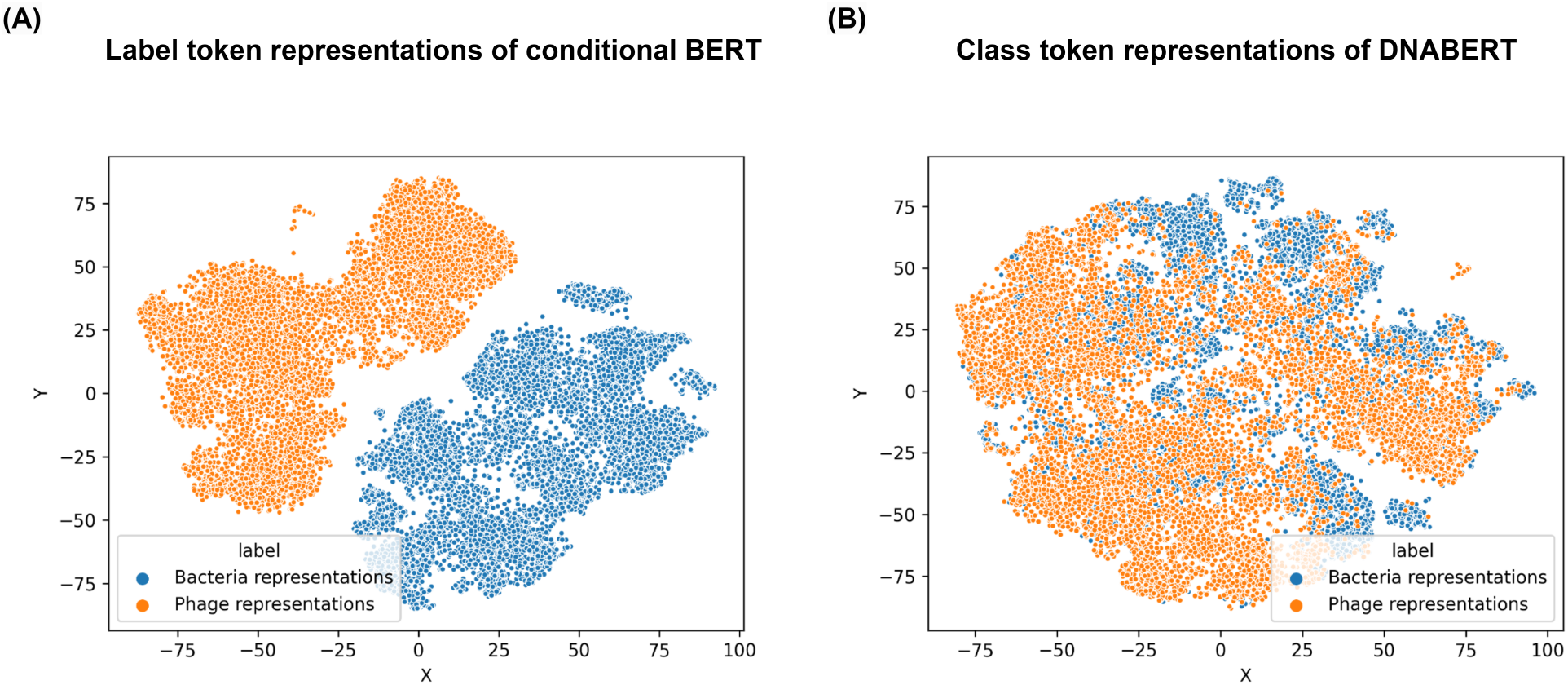
The t-SNE plot of class token representations generated from conditional BERT and DNABERT. (A) The class token representations from Conditional BERT are well-clustered distribution and distinct for bacterial and phage sequences This well-clustered distribution in conditional BERT highlights its ability to learn label-specific contextual information. (B) The class token representations of DNABERT are mixed with bacterial and phage data since it uses the same class token for all sequences.

In contrast, the t-SNE plot for conditional BERT shows well-clustered representations based on the bacterial and phage samples. During pre-training, bacterial and phage samples were attached with different label class tokens: [BAC] for bacteria and [PHA] for phages. The conditional BERT model effectively learns separate contextual representations for different labels during pre-training, forming distinct class token representations for bacteria and phages that are far apart.

This attachment allowed conditional BERT to learn features specific to each label class, resulting in distinct label class token representations for bacterial and phage sequences. This distinction in label class token representations has the potential to be advantageous for applications requiring the distribution of bacteria and phages as prior information, utilizing the model to various downstream tasks.

### 2.2 PharaCon

Previously, conditional BERT models with similar structures have been used for data augmentation in natural language processing (Wu *et al*., 2019). Few of them, however, are directly used for downstream classification tasks. In this study, we propose a solution to use the conditional BERT for classification tasks, predicting the same label classes used in pre-training.

Here, we propose a method based on conditional BERT to identify phages from mixed bacterial and phage sequences, called PharaCon. This section introduces our novel design for fine-tuning PharaCon for the phage identification task.

#### 2.2.1 Paradox samples

In our pre-training process, we employed two types of tokenization based on the sequence type. For bacterial sequences, we appended the label token [BAC], and for phage sequences, we appended the label token [PHA]. This approach allows us to pre-train the sequences with their respective ground truth labels. However, during applications, the ground truth is unknown, resulting in a 50% chance of appending an incorrect label token to the sequence. We refer to this scenario, where an incorrect label is appended, as a “**paradox**” because it deviates from the normal settings established during pre-training. This paradoxical scenario was not included in our pre-training process, as the model was trained to learn contextual information with the correct attachment (the segment tokens attached with their corresponding true labels). Therefore, it is necessary to design a model capable of understanding the relationship between the attached label token and the segment tokens, and distinguishing whether the attached label token is correct. The model should identify the paradox samples from those attached with their ground truth labels.

To address this, we implemented a special augmentation process called paradox augment during fine-tuning and testing. During tokenization and label attachment, each sample is tokenized twice, each time appending a different label token. Bacterial samples labeled with [BAC] are classified as bacteria, phage samples labeled with [PHA] are classified as phages, and metagenomic samples with incorrect label tokens are classified as paradox. This paradox augment process enables conditional BERT to classify bacterial and phage samples smoothly.

#### 2.2.2 Fine-tuning

Our paradox augment process requires designing a model capable of understanding and differentiating the relationship between the attached label token and the segment tokens. To achieve this, we developed a conditional BERT+BiLSTM model to train on the paradox-augmented data within a multi-label classification scheme (see Figure 2B).

During fine-tuning, each segment in the training set is tokenized using the paradox augment strategy. This means each sample is tokenized twice, each time with a different label token attached. These paradox-augmented samples are then fed into the conditional BERT, initialized with the pre-trained weights, to generate their representations.

The following BiLSTM layer processes these representations, learning the relationships between the label representations and the segment representations. This relationship helps the model to differentiate between correctly and incorrectly labeled samples. Finally, a multi-layer perceptron (MLP) is used to make the final prediction based on these learned features.

We named this model framework PharaCon. This model framework not only reinforces the representations of correctly labeled samples learned during pre-training but also helps distinguish them from incorrectly labeled samples, which were not seen during pre-training.

#### 2.2.3 Prediction

Figure 2C illustrates the prediction process of PharaCon. Given a test metagenomic sequence, it is first split into 500 bp-long segments. Each segment is then tokenized with both label tokens, following the paradox augment process, generating two sets of tokens for each segment. PharaCon processes these sets of tokens and generates label class scores for each set. Since each segment generates two sets of tokens, and each set of tokens is predicted with three scores, there are nine combinations of results for each segment. Developing a rule for each of these nine combinations would be overly complex and not robust.

To address this, we propose a label aggregation strategy to simplify the process by aggregating the three-label scores into binarized scores with only bacteria and phage scores. The binarized scores can then be averaged together, regardless of which label token is attached.

Figure 2D details the label aggregation strategy. For a set of tokens, when the label token [PHA] is attached, predicting “paradox” carries the same meaning as predicting bacteria, since “paradox” indicates the attached label is not the ground truth label. Therefore, we add the score of bacteria to the score of paradox. This same logic applies when the label token [BAC] is attached; we add the score of phage to the score of paradox. This approach allows us to convert the three-label scores into binarized scores, containing only bacteria and phage scores. The binarized scores for each set of tokens are then averaged to obtain the final scores for the test sample. To compare with other methods, if the phage score is above 0.5, we predict the test sample as a phage; otherwise, it is predicted as a bacterium. This label aggregation strategy is simple yet effective for handling all possible results, enabling PharaCon to be compared effectively with other methods.

## 3 Numerical experiments

In this study, we designed two experiments to evaluate the performance of PharaCon compared to other selected existing methods. We selected several representative methods: VirSorter2 (Guo *et al*., 2021) and VIBRANT (Kieft *et al*., 2020) for alignment-based approaches, DeepVirFinder (CNN-based) (Ren *et al*., 2020), Seeker (LSTM-based) (Auslander *et al*., 2020), MetaPhaPred (dna2vec+attention mechanism) (Ma *et al*., 2023), and INHERIT (dual Transformer encoders) (Bai *et al*., 2022) for learning-based approaches. To compare these methods with PharaCon from different perspectives, we constructed two datasets: a simulated sequence dataset and a real metagenomic contig dataset. We evaluated the performance of the methods using the following metrics in the experiments: Area Under the Receiver Operating Characteristic Curve (AUROC), Area Under the Precision-Recall Curve (AUPRC), Accuracy, F1 score, and Matthews Correlation Coefficient (MCC). We did not evaluate AUROC and AUPRC for alignment-based methods to avoid unfair comparisons, as they do not generate scores for all test samples. The experimental setup for each method is described in Supplementary materials.

### 3.1 Pre-training dataset and settings

There are two methods, INHERIT and PharaCon, that require pre-training. We used the same datasets as those used when pre-training INHERIT. For bacterial samples, we used the ncbi-genome-download tool (available at https://github.com/kblin/ncbi-genome-download) to download complete bacterial reference genomes published before June 1, 2021, from the NCBI FTP server. Due to physical memory limitations, we randomly sampled 4,124 bacterial genomes, generating 15,975,346 segments.

For phage samples, we directly downloaded and cleaned the sequences longer than 500 bp from NCBI using the keyword “phage” for entries published before June 22, 2021. We also included samples used in Seeker (Auslander *et al*., 2020) and VIBRANT (Kieft *et al*., 2020). This dataset ultimately contains 26,920 phage sequences, generating 1,750,662 segments.

We pre-trained the conditional BERT model with mixed bacteria and phage datasets using masked language modeling with label constraints. We set the learning rate to 4e-4 and the total batch size to 2,048. There were 10,000 warm-up steps, and the weight decay rate was 0.01.

For INHERIT, we directly used the pre-trained weights of the bacteria and phage models available at https://github.com/Celestial-Bai/INHERIT.

### 3.2 Evaluation on simulated sequence dataset

#### 3.2.1 Dataset Information

We built the simulated sequence dataset based on the information provided by MetaPhaPred (Ma *et al*., 2023). Using their guidelines, we downloaded bacterial and phage sequences released before October 31, 2022, from the National Center for Biotechnology Information (NCBI) database. Following their settings, we used 11,505 bacterial and 10,639 phage sequences released before January 1, 2020, as the training sets. However, we split the sequences into fixed 1000 bp-long segments, since most learning-based methods (DeepVirFinder, Seeker, INHERIT, PharaCon) are trained with 500 bp or 1000 bp segments as default settings. We randomly sampled 100,000 bacterial and phage segments to build the training set. If the input length of a model should be set at 500 bp, we split those 1000 bp segments into 500 bp fragments.

To prevent overlap with the pre-training set, we selected 1,302 bacterial and 327 phage sequences released between July 1, 2021, and October 31, 2022, for testing. We cleaned the redundant sequences in the test set by ensuring that if there were identical sequences with different accessions, only one was selected. Since the segments in the training sets are relatively short, we designed a data cleaning strategy specifically suited for short sequences. If a sequence in our selected set had a region overlapping with the segments in the pre-training set or training set detected by BLASTN (McGinnis and Madden, 2004) with 100% identity, we excluded it from the final test set.

Ultimately, we cleaned 434 bacterial accessions from 61,632 accessions in the 1,302 bacterial samples and 241 phage samples. We randomly split the remaining bacterial and phage samples into segments ranging from 500 to 10,000 bp in length. For phage samples, this generated 1,302 segments. To balance the positive and negative samples, we randomly sampled 1,302 segments from the generated bacterial segments.

#### 3.2.2 Experimental results

Table 1 shows the overall performance of PharaCon and other representative methods in simulated sequence dataset. PharaCon emerged as the top performer across all metrics, achieving the highest AUROC (0.9974), AUPRC (0.9964), accuracy (0.9830), F1 score (0.9831), and MCC (0.9661). This indicated that PharaCon was highly effective in distinguishing between phages and bacteria, maintaining a high balance between precision and recall.

**Table 1.** Performance of PharaCon, INHERIT, VIBRANT, VirSorter2, Seeker, DeepVirFinder, and MetaPhaPred in the simulated sequence dataset. PharaCon reached the highest performance with MCC at 0.9661.

| Method | AUROC | AUPRC | Accuracy | F1 score | MCC |
| --- | --- | --- | --- | --- | --- |
| Learning-based methods |  |  |  |  |  |
| PharaCon | **0.9974** | **0.9964** | **0.9830** | **0.9831** | **0.9661** |
| INHERIT | 0.9965 | 0.9956 | 0.9690 | 0.9683 | 0.9388 |
| MetaPhaPred | 0.9921 | 0.9934 | 0.9690 | 0.9693 | 0.9382 |
| DeepVirFinder | 0.9856 | 0.9868 | 0.9399 | 0.9401 | 0.8798 |
| Seeker | 0.8741 | 0.8769 | 0.8081 | 0.8042 | 0.6168 |
| Traditional pre-training strategy |  |  |  |  |  |
| DNABERT+MLP | 0.9939 | 0.9946 | 0.9278 | 0.9225 | 0.8637 |
| DNABERT+BiLSTM | 0.9947 | 0.9954 | 0.9356 | 0.9313 | 0.8778 |
| Alignment-based methods |  |  |  |  |  |
| VIBRANT | - | - | 0.9598 | 0.9606 | 0.9210 |
| VirSorter2 | - | - | 0.9351 | 0.9326 | 0.8725 |

PharaCon demonstrated a slight edge over INHERIT, highlighting the advantages of conditional pre-training. Specifically, when pre-training the same bacterial and phage samples, using a single BERT model with a conditional pre-training strategy can achieve better performance than using two separate BERT models pre-training bacteria and phages independently. This improvement is attributed to the ability of the conditional BERT to incorporate label-specific contextual information during pre-training, allowing the model to learn more distinct and relevant features for each label. As a result, PharaCon benefits from a more integrated and efficient representation of the data, leading to better performance in phage identification.

Figure 3 presents the confusion matrices of PharaCon, depicting results based on three-label scores and label-aggregated scores. In the three-label confusion matrix, PharaCon only misclassifies normal bacterial and phage samples (i.e., samples with correct labels) as paradox. Notably, all 17 paradox samples misclassified as bacteria were originally attached with the [BAC] label token, and all 17 misclassified as phages were attached with the [PHA] label token. In this dataset, PharaCon consistently predicts the sample as either the attached label or paradox, suggesting that the label token serves as a strong guiding constraint. This constraint enables PharaCon to discern whether the segment features align with the label token features and match the label-specific representations acquired from pre-training. Furthermore, PharaCon's accuracy in predicting paradox samples is comparable to, or even slightly better than, its accuracy for normal samples. After applying label aggregation, the prediction of paradox samples enhanced PharaCon's overall accuracy, resulting in predictions that are marginally more accurate than those for normal samples. This underscores the significance of paradox samples in the fine-tuning and prediction process and demonstrates the robustness of our label aggregation strategy.

To evaluate the performance of the individual methods for samples of varying lengths within the test set, we divided the samples into four sub-test sets based on their lengths' quartiles: 500-3,000 bp, 3,000-5,000 bp, 5,000-7,500 bp, and 7,500-10,000 bp. We then calculated the performance of each method within these four subsets. Figure 4 shows the MCC score of all methods across different length ranges. From the Figure 4, PharaCon consistently outperforms other methods across all length subsets, demonstrating its robustness and effectiveness in identifying phages from bacterial sequences.

**Fig. 4:**
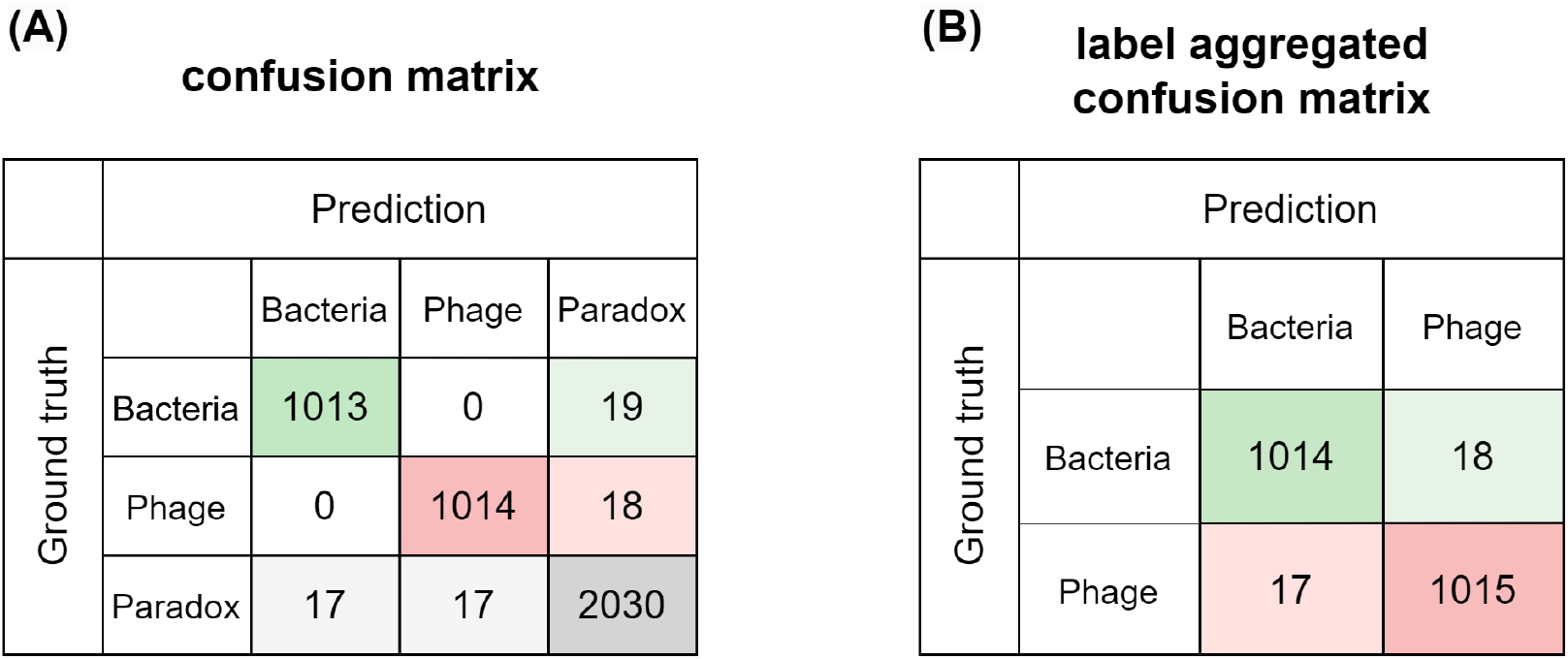
(A) confusion matrix of three label scores generated from PharaCon shows label tokens as strong contraints. (B) Compared the label aggregated confusion matrix of PharaCon with the three-label score confusion matrix, paradox samples can help model better distinguish bacterial and phage samples to some extent.

**Fig. 5:**
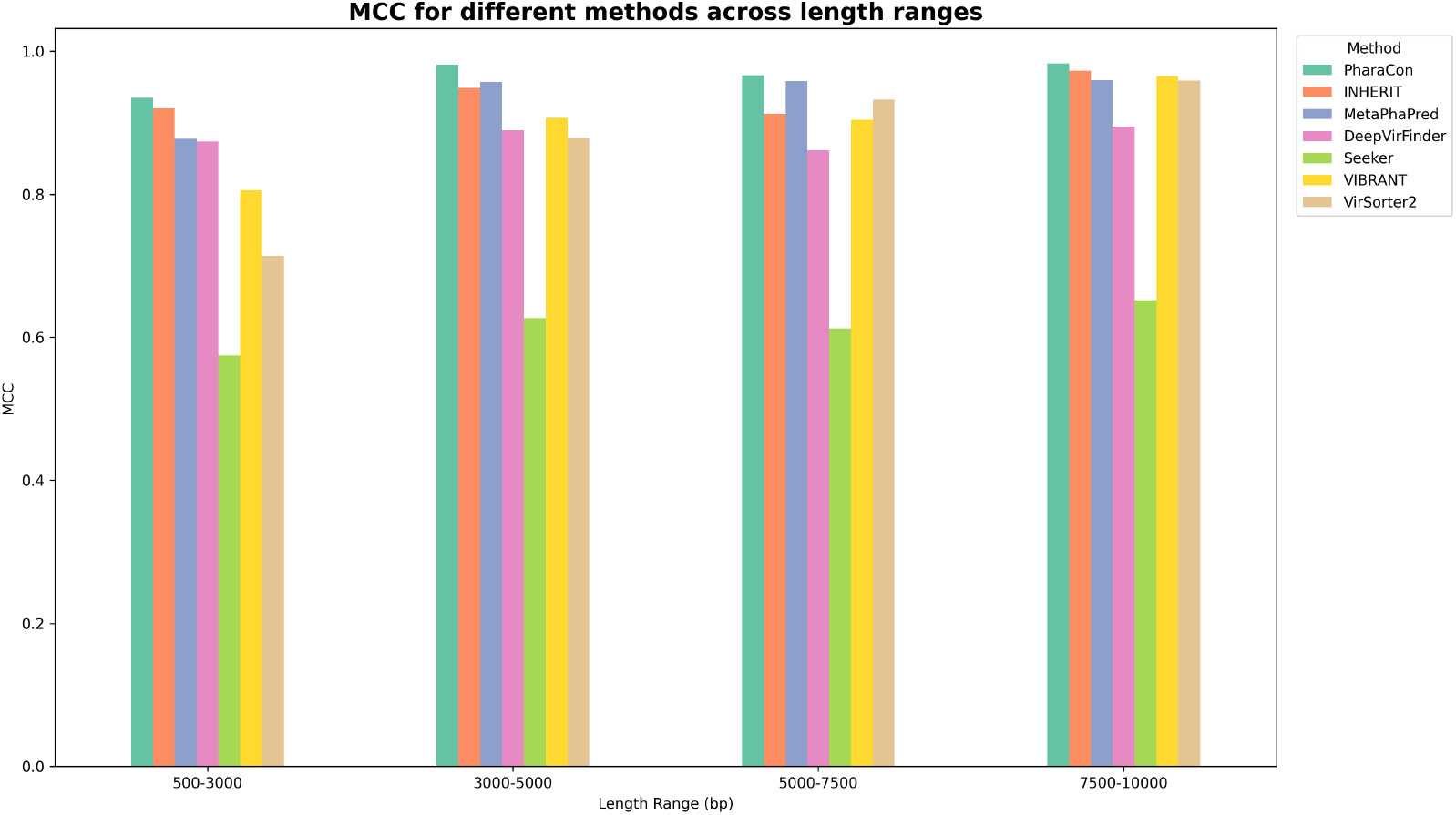
MCC score of PharaCon with other existing methods (Alignment-based: VIBRANT and VirSorter2, and Learning-based: Seeker, DeepVirFinder and MetaPhaPred) in the simulated sequence dataset. We split the dataset into four subsets based on the length's qualtiles. PharaCon achieved the highest MCC in all length ranges.

The alignment-based methods, VIBRANT and VirSorter2, demonstrated varying levels of performance across different sequence lengths, with steady improvements as the sequence lengths increased. This suggests that alignment-based methods are more effective with longer sequences. Additionally, the samples in the test set were submitted after July 1, 2021, which is later than the release of VIBRANT and VirSorter2. This temporal discrepancy may result in less overlap between these test samples and the sequences in the databases used by these alignment-based methods, thereby reducing the advantage of sequence alignment. This lack of overlap could be a significant factor contributing to the relatively poor performance of the alignment-based approaches in this experiment.

Overall, PharaCon consistently achieved the highest performance across all sequence lengths in the simulated sequence dataset, demonstrating robustness and effectiveness with MCC values ranging from 0.9350 to 0.9829. The performance of the methods in the four subsets evaluated with other metrics is shown in the Supplementary Materials.

#### 3.2.3 Comparison with traditional pre-training strategy

We also pre-trained a DNABERT model in this experiment, with the same mixed data and hyperparameters as PharaCon, but using the traditional tokenization strategy and masked language modeling task. We then fine-tuned DNABERT using different classifiers. Initially, we used an MLP classifier, following the traditional sequence classification task. Additionally, we used a BiLSTM classifier with the same structure as used in PharaCon. From the results (see Table 1), neither DNABERT+MLP nor DNABERT+BiLSTM performed better than PharaCon, or even DeepVirFinder. Although PharaCon and DNABERT are both BERT-style frameworks, the differences in pre-training and fine-tuning strategies result in a significant performance gap.

This indicates that compared to the traditional strategy of pre-training with mixed sequence data regardless of labels, our label-specific pre-training strategy is clearly more effective. Even when we changed the downstream classifier from MLP to BiLSTM, the improvement was limited, with the MCC increasing from 0.8637 to 0.8778. The contribution of changing the classifier structure is much less significant than the contribution of the novel label-specific pre-training strategy. The performance gap between these models and PharaCon highlights the advantages of pre-training with label constraints and the paradox-augmented fine-tuning strategy employed in PharaCon.

#### 3.2.4 Extended study on migrating data bias

Since we utilized a supervised pre-training task for PharaCon, the information PharaCon learns is derived from both a large pre-training set and a fine-tuning set. PharaCon uses these datasets as sources for acquiring information to learn features during both pre-training and fine-tuning. Compared to methods without pre-training (MetaPhaPred, DeepVirFinder, Seeker, etc.), PharaCon accesses a significantly larger source of information. This disparity may lead to an unfair comparison, as PharaCon's advantage could stem from its access to more extensive datasets. It is worth mentioning that MetaPhaPred uses pre-trained dna2vec (Ng, 2017) as its embedding layer. However, since dna2vec was pre-trained on the human genome and not on the metagenome, we still consider MetaPhaPred as a method that does not use pre-training.

To eliminate the effect of data bias between methods that utilized pre-training and those that did not, we chose to re-train MetaPhaPred using both the pre-training set and the fine-tuning set while keeping the default hyperparameter settings. The pre-training set includes a large number of sequences, but it contains an imbalance between phage and bacterial samples. To address this, we implemented a down-sampling strategy for bacterial samples to ensure a balanced dataset suitable for training a classifier. First, we split the phage sequences in the pre-training set into 1,000 bp fragments, generating 851,645 segments. Then, we split the bacterial sequences in the pre-training set into 1,000 bp fragments and randomly sampled 851,645 segments from them. We combined these fragments with the samples in the fine-tuning set to re-train MetaPhaPred. By re-training MetaPhaPred with this extended dataset, we aimed to provide a fair comparison with methods that utilized pre-training, ensuring that the performance differences were not solely due to the amount of training data.

After testing with the same dataset, we observed that the MCC of MetaPhaPred improved from 0.9382 to 0.9535, indicating the advantage of increasing the amount of training data. The performance of the re-trained MetaPhaPred with the extended dataset surpassed that of INHERIT but did not exceed PharaCon. This suggests that PharaCon remains the better-performing method even with similar amounts of information access.

It should be emphasized that training models on such extended datasets is not the usual practice. Typically, models are fine-tuned on datasets that are much smaller than the pre-training set to save computational costs. Additionally, the data used in the pre-training set is not filtered as rigorously as the fine-tuning set, and its quality is not as high. This may hinder models with smaller parameters from effectively extracting and learning features from a large dataset. For example, we similarly re-trained Seeker and found that its MCC decreased from 0.6168 to 0.4985. This suggests that in a real-world scenario, it is not always beneficial to train with an excessively large dataset to improve a method's capability. Detailed results are provided in the Supplementary Materials.

### 3.3 Evaluation on real metagenomic contig dataset

#### 3.3.1 Dataset Information

In this section, we focus on evaluating PharaCon and comparing it with other methods using metagenomic contigs from existing prokaryote and viral databases to simulate real-world application scenarios. We constructed datasets based on two phage metagenomic databases: the Gut Virome Database (GVD) (Gregory *et al*., 2020) and the Cenote Human Virome Database (CHVD) (Tisza and Buck, 2021), as well as a bacterial metagenomic database called the Unified Human Gastrointestinal Genome (UHGG) (Almeida *et al*., 2021). We also constructed a training set for learning-based methods, as the feature distribution between reference genomes and real metagenomic contigs is different (see Supplementary Materials). Considering computational resources, we used the contigs published before 2019 in the GVD database for the training set. We randomly selected 20% of them as validation samples.

Since learning-based methods operate and predict at the segment level, we randomly selected bacterial sequences from UHGG to build the training and validation samples to match the total length of phage sequences in the training and validation sets, respectively. This ensured a balance between bacterial and phage segments. In summary, there are 12,949 bacterial contigs and 12,585 phage contigs in the training set, and 3,716 bacterial contigs and 3,142 phage contigs in the validation set.

The phage samples in the test set were obtained from the CHVD database. We removed phages with cognate sequences in the training set from the CHVD database. The bacterial contigs in the test set were also randomly sampled from UHGG and did not overlap with those in the training set. The number of bacterial and phage contigs was equal in the test set. In summary, there are 36,012 bacterial contigs and 36,012 phage contigs in the test set.

#### 3.3.2 Experimental results

According to the results (see Table 2), the alignment-based methods, VirSorter2 and VIBRANT, exhibited strong performance on the real metagenome dataset. VirSorter2 outperformed all other methods, both learning-based and alignment-based. Among the learning-based methods, PharaCon led in all metrics, showcasing its capability in identifying phages. The performance trend of learning-based methods in the real metagenomic contig dataset is consistent with the trend observed in the simulated sequence dataset, further highlighting the advantages of the label-specific pre-training strategy with conditional BERT. By leveraging label-specific information during pre-training and employing a paradox-augmented fine-tuning strategy, PharaCon effectively enhances identification ability and demonstrates advantages over other learning-based methods.

**Table 2.** Performance of PharaCon, INHERIT, VIBRANT, VirSorter2, Seeker, DeepVirFinder, and MetaPhaPred in the real metagenomic contig dataset. PharaCon is the best performing learning-based methods, while VirSorter2 reached the best performance.

| Method | AUROC | AUPRC | Accuracy | F1 score | MCC |
| --- | --- | --- | --- | --- | --- |
| Learning-based methods |  |  |  |  |  |
| PharaCon | **0.9719** | **0.9638** | 0.9128 | 0.9097 | 0.8274 |
| INHERIT | 0.9592 | 0.9439 | 0.8985 | 0.8965 | 0.7975 |
| MetaPhaPred | 0.9371 | 0.9217 | 0.8637 | 0.8693 | 0.7301 |
| DeepVirFinder | 0.9155 | 0.9025 | 0.8082 | 0.7859 | 0.6301 |
| Seeker | 0.843 | 0.8105 | 0.7648 | 0.7698 | 0.5301 |
| Alignment-based methods |  |  |  |  |  |
| VIBRANT | - | - | 0.9155 | 0.9199 | 0.8383 |
| VirSorter2 | - | - | **0.9635** | **0.9639** | **0.9272** |

However, in this dataset, no learning-based method performed better than the alignment-based methods. While PharaCon performed comparably to VIBRANT, it still had a 0.14% lower MCC than VIBRANT. The main reason why alignment-based methods performed better than all learning-based methods could be the identification process of viral contigs in the CHVD database. The viral contigs in the CHVD database were all identified by Cenote-Taker 2 (Tisza *et al*., 2021), a database-based method that includes profile Hidden Markov Models from the Pfam database (Mistry *et al*., 2021). These models are also referenced by VIBRANT and VirSorter2. The high overlap between the profile Hidden Markov Models used to identify the viral contigs in CHVD and those referenced by VIBRANT and VirSorter2 may account for the high precision in VIBRANT (0.9800) and the high recall in VirSorter2 (0.9754).

We also tested the total prediction time for all methods. PharaCon was slower than other learning-based methods except for INHERIT, but the total running time was still acceptable. It demonstrated better performance while running in less time than INHERIT in our experiments. Although VirSorter2 performed the best, it took more than 15,040 minutes to finish predicting all the contigs on our device (see Table 3), making it the slowest among the methods we evaluated. In contrast, PharaCon required approximately 10% of the time that VirSorter2 took to complete the prediction, which is about 85% faster than VIBRANT. PharaCon can achieve a performance level similar to VIBRANT while requiring much less time than database-based methods, proving its effectiveness and efficiency.

**Table 3.** The total prediction time (minute) for PharaCon and other existing methods in the metagenomic contigs dataset. PharaCon ran slower than other learning-based methods except INHERIT, while much faster than database-based methods.

| Method | Total prediction time |
| --- | --- |
| PharaCon | 1,506 |
| INHERIT | 1,744 |
| VIBRANT | 10,076 |
| VirSorter2 | 15,040 |
| DeepVirFinder | 428 |
| Seeker | 77 |
| MetaPhaPred | 408 |

## 4 Discussion

In this paper, we mainly focused on applying conditional BERT to phage identification, introducing PharaCon. The underlying principles could potentially be extended to other genomic classification challenges. The approach of leveraging label constraints during pre-training and the novel fine-tuning strategy may hold broader applicability. We hope that future research will explore the application of conditional BERT frameworks to other genomic classification tasks. Testing the conditional BERT and the novel pre-train-fine-tune paradigm across a variety of genomic classification tasks will help determine its robustness and versatility, which could provide valuable insights into its broader applicability.

Furthermore, the current work formulated phage identification as a binary classification problem, distinguishing bacterial from phage sequences. However, genomic data in real-world scenarios often exhibits multi-label characteristics, wherein sequences may simultaneously belong to multiple categories. Consequently, there is a pressing need for models capable of handling multi-label classification tasks. We hope that future research will investigate on extending the conditional BERT framework and the proposed fine-tuning scheme to support multi-label classification, thereby enhancing their utility and effectiveness in more intricate genomic analyses.

## 5 Conclusion

In this study, we proposed a novel conditional BERT model for pre-training labeled data, incorporating label constraints during pre-training with modified language modeling tasks. Additionally, we applied conditional BERT to the phage identification task with a paradox-augmented fine-tuning strategy, introducing PharaCon. We tested PharaCon against other selected existing methods on both simulated sequence datasets and real metagenomic contig datasets. PharaCon outperformed other learning-based methods across multiple metrics in both datasets and achieved the best MCC score of 0.9661 in the simulated sequence dataset among all methods. In the real metagenomic contig dataset, it was the only learning-based method to achieve performance competitive with alignment-based methods like VIBRANT, while significantly reducing prediction time. The innovative use of label constraints during pre-training and the paradox-augmented fine-tuning process contributed to its enhanced accuracy and efficiency, making PharaCon an effective method for phage identification in metagenomic sequences.

## Acknowledgements

The super-computing resource was provided by the Human Genome Center, Institute of Medical Science, The University of Tokyo (https://gc.hgc.jp/en/).

