## Supplementary material for "PharaCon: A new framework for identifying bacteriophages via conditional representation learning": Supplemtary materials

**Supplementary Materials**

**Table S1	Experimental setups**

We made the following settings for the selected methods based on their default configurations to evaluate and compare them as fairly as possible. These tables summarize the configurations for each method used in the experiments for both datasets.

**Simulated sequence dataset**

| **Method** | **Training Length** | **Epochs** | **Batch Size** | **Learning Rate** | **Training Device** | **Additional Settings** |
| --- | --- | --- | --- | --- | --- | --- |
| Learning-based methods | | | | | | |
| PharaCon | 500 bp | 10 | 64 | 1e-5 | NVIDIA A100 | - |
| INHERIT | 500 bp | 10 | 64 | 1e-5 | NVIDIA A100 | - |
| MetaPhaPred | 500 bp, 1,000 bp | 15 | 256 | 1e-3 | NVIDIA A100 | Separate models for sequences shorter and longer than 800 bp. |
| DeepVirFinder | 500 bp, 1,000 bp | 30 | - | 1e-3 | NVIDIA A100 | Separate models for sequences shorter and longer than 1,000 bp |
| Seeker | 1,000 bp | 100 | 27 | - | NVIDIA A100 | - |
| Database-based methods | | | | | | |
| VIBRANT | - | - | - | - | Intel Xeon Gold 6154 | Version 1.2.1; default command from GitHub; excluded segments < 1,000 bp or with < 4 ORFs. |
| VirSorter2 | - | - | - | - | Intel Xeon Gold 6154 | Version 2.1; considered a segment as phage if predicted whole or part as dsDNAphage or ssDNA. |

**Metagenomic contig dataset**

| **Method** | **Batch Size** | **Learning Rate** | **Epochs** | **Training Device** | **Additional Settings** |
| --- | --- | --- | --- | --- | --- |
| Learning-based methods | | | | | |
| PharaCon | 64 | 1e-5 | - | NVIDIA A100 | Checkpoint with best validation loss; validation loss did not decrease for 5 epochs. |
| INHERIT | 64 | 1e-5 | - | NVIDIA A100 | Checkpoint with best validation accuracy; validation accuracy did not increase for 3 epochs. |
| MetaPhaPred | 256 | 1e-3 | 15 | NVIDIA A100 | Three models for sequences shorter than 400 bp, 401-800 bp, and longer than 800 bp. |
| Seeker | 27 | - | 100 | NVIDIA A100 | - |
| DeepVirFinder | - | 1e-3 | 30 | NVIDIA A100 | Four models for sequences shorter than 300 bp, 300-500 bp, 500-1,000 bp, and longer than 1,000 bp. |
| Database-based methods | | | | | |
| VIBRANT | - | - | - | Intel Xeon Gold 6154 | Same settings as the simulated sequence dataset. |
| VirSorter2 | - | - | - | Intel Xeon Gold 6154 | Same settings as the simulated sequence dataset. |

**Table S2	The performance of each method on simulated sequence dataset with different length ranges**

|  | AUROC | AUPRC | Accuracy | F1 | MCC |
| --- | --- | --- | --- | --- | --- |
| 500-3000: bacteria: 251 phages: 272 | | | | | |
| PharaCon | **0.9935** | **0.9923** | **0.9675** | **0.9690** | **0.9350** |
| INHERIT | 0.9916 | 0.9909 | 0.9598 | 0.9607 | 0.9202 |
| MPP | 0.9788 | 0.9834 | 0.9388 | 0.9420 | 0.8777 |
| DeepVirFinder | 0.9837 | 0.9866 | 0.9369 | 0.9401 | 0.8738 |
| Seeker | 0.8535 | 0.8652 | 0.7859 | 0.7854 | 0.5745 |
| VIBRANT | 0.9394 | 0.9690 | 0.9098 | 0.9355 | 0.8059 |
| VirSorter2 | 0.8512 | 0.8399 | 0.8470 | 0.8354 | 0.7138 |
| 3000-5000: bacteria: 209 phages: 209 | | | | | |
| PharaCon | **0.9996** | **0.9996** | **0.9904** | **0.9904** | **0.9809** |
| INHERIT | 0.9996 | 0.9996 | 0.9737 | 0.9730 | 0.9487 |
| MPP | 0.9957 | 0.9958 | 0.9785 | 0.9785 | 0.9569 |
| DeepVirFinder | 0.9871 | 0.9875 | 0.9448 | 0.9448 | 0.8897 |
| Seeker | 0.8837 | 0.8814 | 0.8134 | 0.8116 | 0.6269 |
| VIBRANT | 0.9555 | 0.9570 | 0.9521 | 0.9543 | 0.9073 |
| VirSorter2 | 0.9378 | 0.9286 | 0.9378 | 0.9350 | 0.8789 |
| 5000-7500: bacteria: 255 phages: 279 | | | | | |
| PharaCon | **0.9978** | **0.9979** | **0.9831** | **0.9838** | **0.9662** |
| INHERIT | 0.9981 | 0.9982 | 0.9551 | 0.9556 | 0.9123 |
| MPP | 0.9975 | 0.9976 | 0.9794 | 0.9804 | 0.9588 |
| DeepVirFinder | 0.9833 | 0.9855 | 0.9307 | 0.9326 | 0.8618 |
| Seeker | 0.8762 | 0.8931 | 0.8052 | 0.8074 | 0.6123 |
| VIBRANT | 0.9528 | 0.9470 | 0.9514 | 0.9526 | 0.9043 |
| VirSorter2 | 0.9666 | 0.9567 | 0.9663 | 0.9675 | 0.9326 |
| 7500-10000: bacteria: 317 phages: 272 | | | | | |
| PharaCon | **0.9996** | **0.9996** | **0.9915** | **0.9908** | **0.9829** |
| INHERIT | 0.9993 | 0.9992 | 0.9864 | 0.9852 | 0.9727 |
| MPP | 0.9986 | 0.9985 | 0.9796 | 0.9783 | 0.9595 |
| DeepVirFinder | 0.9906 | 0.9904 | 0.9474 | 0.9439 | 0.8948 |
| Seeker | 0.8884 | 0.8763 | 0.8268 | 0.8132 | 0.6518 |
| VIBRANT | 0.9825 | 0.9733 | 0.9829 | 0.9813 | 0.9656 |
| VirSorter2 | 0.9798 | 0.9650 | 0.9796 | 0.9780 | 0.9591 |

**Table S3	The performance of MetaPhaPred re-trained with both pre-training data and fine-tuning data (noted as MetaPhaPred *)**

|  | AUROC | AUPRC | Accuracy | F1 | MCC |
| --- | --- | --- | --- | --- | --- |
| MetaPhaPred* | 0.9961 | 0.9966 | 0.9767 | 0.9768 | 0.9535 |
| MetaPhaPred | 0.9921 | 0.9934 | 0.9690 | 0.9693 | 0.9382 |

**Table S4	The performance of Seeker re-trained with both pre-training data and fine-tuning data (noted as Seeker*)**

|  | AUROC | AUPRC | Accuracy | F1 | MCC |
| --- | --- | --- | --- | --- | --- |
| Seeker* | 0.8794 | 0.8797 | 0.7219 | 0.7735 | 0.4985 |
| Seeker | 0.8741 | 0.8769 | 0.8081 | 0.8042 | 0.6168 |
